# Lost in the Forest: Encoding Categorical Variables and the Absent Levels Problem

**DOI:** 10.1101/2022.09.12.507676

**Authors:** Helen L. Smith, Patrick J. Biggs, Nigel P. French, Adam N.H. Smith, Jonathan C. Marshall

## Abstract

Levels of a predictor variable that are absent when a classification tree is grown can not be subject to an explicit splitting rule. This is an issue if these absent levels then present in a new observation for prediction. To date, there remains no satisfactory solution for absent levels in random forest models. Unlike missing data, absent levels are fully observed and known. Ordinal encoding of predictors allows absent levels to be integrated and used for prediction. Using a case study on source attribution of *Campylobacter* species using whole genome sequencing (WGS) data as predictors, we examine how target-agnostic *versus* target-based encoding of predictor variables with absent levels affects the accuracy of random forest models. We show that a target-based encoding approach using class probabilities, with absent levels designated the highest rank, is systematically biased, and that this bias is resolved by encoding absent levels according to the *a priori* hypothesis of equal class probability. We present a novel method of ordinal encoding predictors *via* principal coordinates analysis (PCO) which capitalizes on the similarity between pairs of predictor levels. Absent levels are encoded according to their similarity to each of the other levels in the training data. We show that the PCO-encoding method performs at least as well as the target-based approach and is not biased.

## Introduction

### 1.1 Random forest

A classification tree is a method of supervised machine learning that predicts a categorical response variable by way of a series of binary decisions. Random forest is an ensemble of classification trees that aggregates the predictions from the individual trees to inform a classification.

### 1.2 The ‘absent levels’ problem

An inherent issue with tree-based predictive models occurs when a level of a categorical predictor variable is absent when a tree is grown, but is present in a new observation for prediction (the ‘absent levels’ problem *sensu* Au, 2018). In a random forest, absent levels can arise due to sampling variability (i.e., the level was absent from the observations that were used to train the model), bagging (i.e., the level was in the training data but absent from the bootstrapped sample used by a particular tree), or partitioning of the data by the tree (i.e., the level was present at the top of the tree but absent from a lower subset created by binary splits). When the algorithm encounters an absent level, there is no immutable *a priori* rule for determining which side of the binary split the observation should go. When this happens, the observation is effectively ‘lost in the forest’.

Missing data heuristics allow the random forest algorithm to proceed with an absent level. Methods include stopping an affected observation from proceeding down the tree (Therneau, Atkinson, & Ripley, 2022), using a surrogate decision rule that mimics the original split’s partitioning (i.e. ‘surrogate splits’) (Hothorn & Zeileis, 2015; Therneau et al., 2022), directing all affected observations down the branch with the most training observations (Hothorn & Zeileis, 2015), directing all affected observations down both branches simultaneously but weighted according to the number of observations from each child node (e.g. distribution-based imputation (DBI)) (Quinlan, 1993; Saar-Tsechansky & Provost, 2007), and randomly directing affected observations down a left or right branch (Hothorn & Zeileis, 2015). The `scikit-learn` Python module’s implementation of random forests (Pedregosa et al., 2011) treats absent levels as missing values. If missing levels are present in the training data then absent levels get assigned to an explicit missing category, otherwise they get mapped to the child node that has the most samples^1^.

It is important to distinguish between absent levels and missing data, however. Unlike missing data, absent levels are fully observed and known. Treating an observation with an absent level as though it were missing data necessitates a loss of information and is not recommended (Ishwaran, Kogalur, Blackstone, & Lauer, 2008). Au (2018) thoroughly investigated the properties of missing data heuristics with random forests and compared them to the naïve heuristic of directing all observations with absent levels down the same branch (i.e., “left” or “right heuristic”) as is implemented in many random forest applications (Liaw & Wiener, 2002; Wright & König, 2019). Au showed that the choice of heuristic can dramatically alter a model’s performance and potentially lead to systematic bias in prediction. Decision tree-based methods are widely used; it is almost certain that a number of these models have been inadvertently affected by the absent levels problem in practice. To date, there remains no compelling solution for dealing with absent levels in random forest models^2^.

### 1.3 Out-of-bag sample

Bootstrap aggregating, or bagging, in random forest sees each individual tree trained on a subset of the observations in the training set generated by subsampling with replacement. Correspondingly, for each tree there is a sample of observations that are not used for training - the out-of-bag (OOB) sample. Aggregating the predictions from the observations in the OOB sample can be used to generate an OOB prediction for each observation; the misclassification rate of OOB predictions for all training observations is the OOB error (Breiman, 2001). Breiman (1996, 2001) claimed that the OOB error alleviates the need for cross-validation or setting aside a separate test set, however, at least for two-class classification problems with numerical predictor variables, this is disputable (Janitza & Hornung, 2018; Mitchell, 2011).

### 1.4 Variable encoding

The key to dealing with absent levels lies in how categorical variables are encoded. The random forest algorithm can, in theory, process categorical variables in their raw state, comparing all 2^*k−*1^*−*1 possible binary splits for a nominal predictor variable with *k* distinct levels. There are, however, significant potential gains in efficiency from imposing an order on a nominal predictor variable. An ordered categorical predictor with *k* levels can be treated the same way as a numerical predictor with *k* unique ordered values. This reduces the number of potential partitions from 2^*k−*1^*−*1 to *k−*1 and the allocation of each level to one side of the binary split is constrained only by whether it is above or below the split point. There are several approaches used to encode, or convert, categorical variables into numerical format for analysis.

Integer encoding (also called label encoding) is the simplest method of encoding. For each categorical variable *X*, with *k* distinct values, the observed levels are mapped to the integers 1 to *k* and new levels, which were not observed during training, are encoded as missing values. A major issue with this method is that despite there being no intrinsic relationship between the levels and the numbers being used to replace them an ordering (1 *< k*) is imposed.

Indicator encoding (also called one-hot encoding) avoids imposing an order on a nominal categorical variable. Each categorical variable *X*, with *k* distinct values, is transformed to *k* binary indicator variables and observations are encoded to indicate the presence (1) or absence (0) of the dichotomous variable. Indicator encoding removes any uncertainty over where to send an observation with an absent level as they can be encoded with a zero vector. However, it can result in the dataset becoming very wide and sparse, which in turn can present computational challenges and inconsistent results (Au, 2018; Cerda, Varoquaux, & Kegl, 2018; Hastie, Tibshirani, & Friedman, 2009; Reilly, Taylor, Fergus, Chalmers, & Thompson, 2022). With indicator encoding, the feature importance of the original variable is distributed among separate binary variables which may cause bias for tree based algorithms as the impurity reduction induced by a single indicator is rarely enough to be selected for splitting. Dummy encoding, i.e. indicator encoding with *k−*1 categories, has similar properties.

Target-based encoding methods differ from integer encoding and indicator encoding in that they incorporate information about the target values associated with a given level. For the case of two-class (binary) classification, ordering a nominal predictor by the proportion of observations with the second response class in each level leads to identical splits in the random forest optimisation as considering all possible 2-partitions of the predictor levels if the encoding is repeated at every split (Breiman, Friedman, Olshen, & Stone, 1984; Fisher, 1958). Two popular software implementations for random forest, the `randomForest` and `ranger` R packages, adopt this optimisation. For multiclass classification, there is no available sorting algorithm that leads to splits which are equivalent to considering all 2^*k−*1^*−*1 possible partitions (Wright & König, 2019). The R package `ranger` (Wright & Ziegler, 2017) offers a target-based encoding method that encodes each predictor variable according to the first principal component of the weighted covariance matrix of class probabilities, following Coppersmith, Hong, and Hosking (1999)^3^. For computational efficiency the encoding of the predictor variables occurs once on the entire dataset prior to bagging. In each of these methods absent levels are encoded with the highest rank, effecting the “right” heuristic.

The method of encoding predictor variables can affect the performance of random forest (Au, 2018; Wright & König, 2019). For categorical features with a small number of levels, target-based encoding has been shown to achieve better results than one-hot encoding and integer encoding (Wright & König, 2019), however as a result of using the target variable, information leakage and overfitting is a concern. By using the probability of the target for encoding, there is information leakage from the target variable to the predictors. Further, if a predictor is encoded prior to splitting into training and testing sets, information from the target variable in the test set will leak to the predictors in the training set by way of the *a priori* encoding, which will impact cross-validation errors. When the encoding occurs prior to bagging (i.e., rather than each subsample undergoing encoding independently) the OOB errors will be similarly affected. A separate test set that is not used to inform the encoding will be a more reliable estimate of model performance.

### 1.5 Encoding of absent levels

In addition to reducing computational complexity, ordinal encoding of predictor variables allows absent levels to be encoded, integrated with existing levels, and subsequently used for prediction, thereby circumventing the absent levels problem. The `randomForest`^4^ and `ranger` R packages encode absent levels with the highest rank (equivalent to integer encoding as *k* + 1), which ensures observations with an absent level will always “go right”, as per the “right” heuristic for missing data. Assigning all observations with absent levels to the same branch will keep the observations together as a collection which can be split further down the tree by another variable. However, this heuristic, when combined with target encoding, leads to systematic bias towards the first response class (Au, 2018). Furthermore, classifications for observations with absent levels can be influenced by interchanging the order of the response classes. Au (2018) therefore argued that observations with absent levels should be assigned randomly to a left or right branch as this reduces the systematic bias in prediction. There has been no documented investigation to date into the properties of this heuristic in the multiclass response case, however.

Here, we examine various methods of dealing with high cardinality, nominal predictor variables in the context of random forest models and the absent levels problem. We detail how target-agnostic *versus* target-based encoding predictor variables with absent levels affects the accuracy of random forest models, and we present two alternate methods for encoding predictor variables and/or absent levels. We examine prediction accuracy of these methods using a case study on source attribution of *Campylobacter* species using whole genome sequencing (WGS) data as predictors. The WGS data generates allele profiles based on unique nucleotide sequences for each gene in the chromosome.

More specifically, we aim to:

i. assess the misclassification rate of multiclass random forest predictions when nominal predictor variables are target encoded and observations with absent levels are sent to the right side of a binary split, using real data from a published source-assigned case-control study;
ii. compare the misclassification rate from (i) *versus* that of predictions when observations with absent levels are sent to a left or a right branch of a split according to the *a priori* hypothesis of equal class probability;
iii. introduce the PCO-encoding method for ordinal encoding categorical predictors that makes use of ancillary information on the levels of predictor variables.

## Methods

### 2.1 Random Forest

For a training set of *N* independent observations on *P* variables, where *x*_*n*_ = (*x*_*n*1_, *x*_*n*2_, …, *x*_*nP*_) is the vector of predictor variables for observation *n* = 1, 2, …, *N*, and *y*_*n*_ is the corresponding response variable, Classification and Regression Tree (CART) is a greedy recursive binary partitioning algorithm that successively partitions data (the parent node) into two smaller subsets (the left and right child nodes).

Each partition is determined based on a decision rule for a single predictor variable to maximise predictive accuracy with respect to the response variable (Breiman et al., 1984). In a random forest, each individual tree is trained on a bootstrap resample of the training data (‘bagging’) using a randomly selected subset of the *P* predictors (‘random subspacing’; Amit & Geman, 1997; Breiman, 1996; Ho, 1998), and is traditionally not “pruned”. A classification can be predicted for a new observation by sending it down each tree according to the decision rules until it arrives at a terminal node, then aggregating the tree predictions and taking the majority vote across the forest. Various control parameters can be set for random forests, including the number of trees, the number of variables randomly selected as splitting candidates, and tree size (Wright & Ziegler, 2017).

### 2.2 Encoding of Categorical Predictor Variables

For multiclass classification problems, we consider three methods for encoding categorical predictor variables as ordered factors or continuous variables:

#### 2.2.1 Correspondence analysis (CA) encoding method

The CA-encoding method is a target-encoding method which performs a scaled correspondence analysis on the contingency table of counts of variable levels by class, following the approximation of Coppersmith, Hong, and Hosking (1999).^5^ Each predictor variable is encoded according to the first principal component of the weighted matrix of class probabilities and absent levels are encoded with a principal component score of infinity. This ensures all observations with an absent level branch as a group and always (i.e., at each node) in the same direction (“go right”) (figure 1, a).

**Fig. 1.**
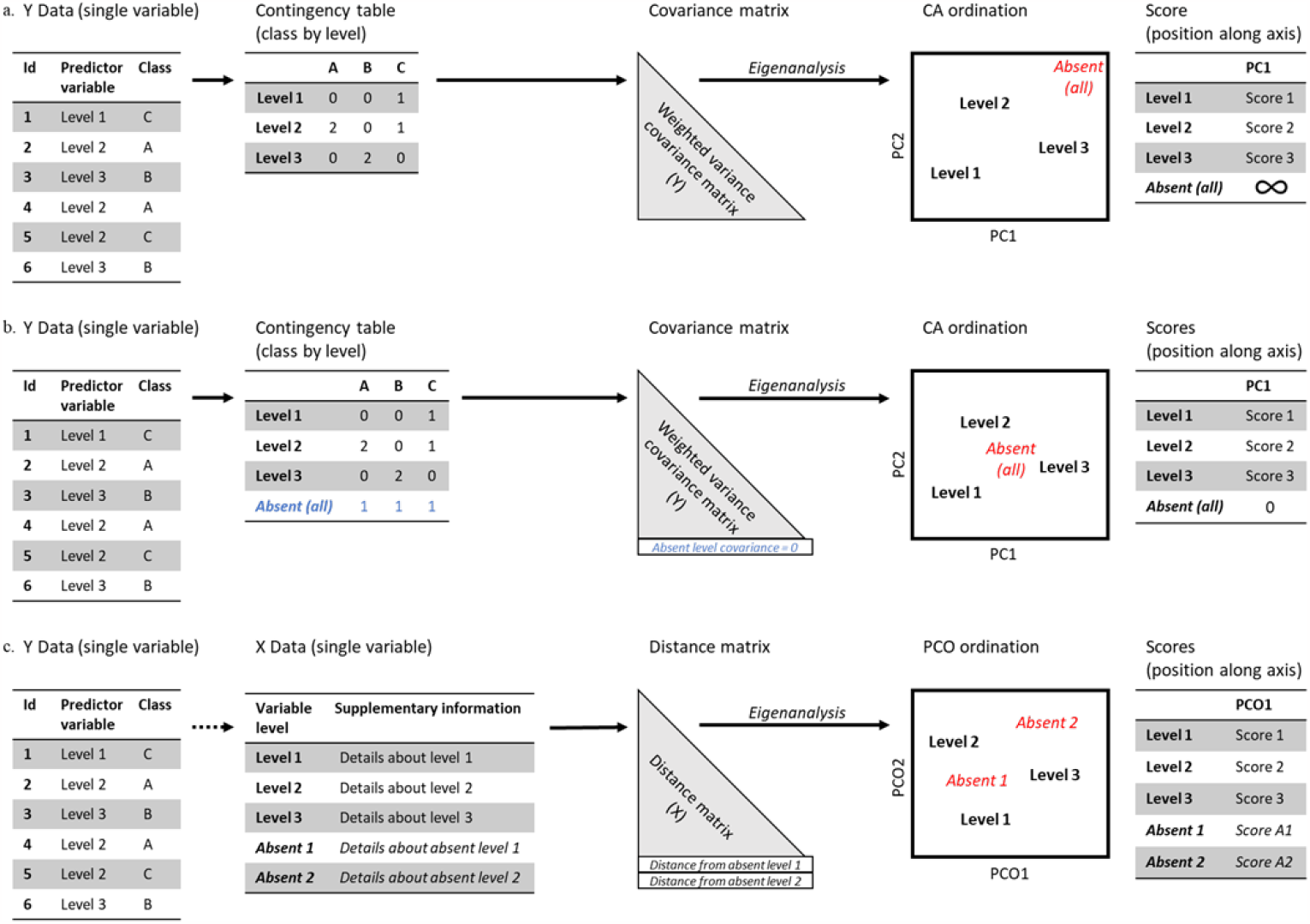
A visual description of the four methods described in this paper (a) CA-encoding method - the levels of each predictor variable are ordered according to the first principal component of the class probabilities and absent levels are assigned a score of infinity; (b) CA-unbiased-encoding method - the levels of each predictor variable are ordered according to the first principal component of the class probabilities and absent levels are assigned a score of zero based on *a priori* equal class probabilities; blue text indicates conceptual information for an absent level; (c) PCO-encoding method - the levels of each predictor variable, including absent levels, are ordered according to their score for the first principal coordinate axis derived from ancillary pair-wise distance information.

#### 2.2.2 CA-unbiased encoding method

The difference between the CA and CA-unbiased methods of encoding lies in the treatment of absent levels. Our novel CA-unbiased-encoding method encodes any absent level with a principal component score of zero (figure 1, b). This aligns with the assumption that any level of the predictor variable that is absent from the training data is *a priori* equally likely in any class and has equal class probabilities of 1*/Y*, where *Y* is the number of classes. Because all absent levels will have equal class probability vectors, they can be combined into a single attribute value (Coppersmith, Hong, & Hosking, 1999). Then, because the class probabilities are not independent of each other, the sum of the principal component coefficients is zero and it follows that the principal component score of an absent level with equal class probabilities will be zero. At some splits, the zero principal component score will fall on the left side of the splitting value and at other splits it will fall on the right side. In the unlikely case of a splitting value being exactly zero, all observations would be sent to the left. To account for this, the scores have a small degree of noise added so that observations will be randomly sent to either the left or right branch with equal probability in the case of a splitting value being exactly zero.

### 2.2.3 Principal coordinates analysis (PCO) encoding method

The PCO-encoding method is a target-agnostic ordinal encoding method which relies on ancillary information on the individual levels of predictor variables. For example, a categorical variable consisting of city names has ancillary information that includes latitude and longitude, as well as population-based information. The PCO-encoding method utilises the ancillary information, rather than the level names *per se*. The choice of ancillary variables used for the distance calculation will depend on how the association between levels should be defined, e.g. geographical *versus* social *versus* economic etc. This will determine the degree of similarity and how an absent level will be treated. In the correspondence analysis methods above, the eigenanalysis step is performed on the weighted level by class contingency table and the score is the coefficient for the corresponding predictor level of the first principal component. In comparison, the eigenanalysis in the PCO-encoding method is performed on a distance matrix of the set of predictor levels extracted from ancillary information on the levels of predictor variables, and the score is the principal component score for the corresponding predictor level for the first principal coordinate (figure 1, c). Our PCO-encoding method relies on ancillary information for each of the predictor variables, independently, in order to generate a set of matrices of dissimilarities. We then apply principal coordinates analysis (PCO) (Gower, 1966) to this distance matrix, yielding a ρ-dimensional ordination of levels in Euclidean space. A single dimension (i.e., only the first principal coordinate) for each variable was chosen to maintain consistency between methods for comparison, however any number of dimensions could potentially be used. Using the method of Gower (1968), a new (absent) level can be interpolated into the ρ-dimensional space by virtue of the interpoint distances between this level and each of the present levels. This then generates a score for each new level, and allows it to branch according to its resemblance to other levels in the training data.

### 2.3 Comparison of Encoding Methods

#### 2.3.1 Source Attribution

The process of assigning cases of human zoonotic infectious diseases to their most likely origin is known as ‘source attribution’. Because of their role in human gastroenteritis, *Campylobacter jejuni* and *C. coli* have been the subject of a large number of source attribution studies using a variety of approaches, including epidemiological methods (Domingues, Pires, Halasa, & Hald, 2012; Pires, Vigre, Makela, & Hald, 2010); comparative risk and exposure assessment (Pintar et al., 2017); expert knowledge elicitation (Hald et al., 2016; Havelaar, Galindo, Kurowicka, & Cooke, 2008); and microbiological methods (Arning, Sheppard, Bayliss, Clifton, & Wilson, 2021; Brinch et al., 2023; Hald, Vose, Wegener, & Koupeev, 2004; Liao, Marshall, Hazelton, & French, 2019; Miller, Marshall, French, & Jewell, 2017; Müllner et al., 2009; Sheppard et al., 2009; Strachan et al., 2009). Microbiological methods of source attribution rely on comparing the phenotypic or genotypic profiles of human cases of infection with those of animal sources. Although many earlier studies have used just a small number of loci (targeted part of a gene in the bacterial chromosome) within the genome (*<* 10), the availability of next generation sequencing has greatly increased the number of loci available for analysis. Models that use allelic-profile data arising from bacterial whole genome sequencing (WGS) have a high number of categorical predictors, which are often subject to the absent levels problem. *Campylobacter* species are genomically very diverse and, although the allelic diversity (i.e., sequence variability within a gene) is inconsistent across the genome, some loci are highly variable (Parkhill et al., 2000; Sheppard & Maiden, 2015). *Campylobacter jejuni* and *C. coli* each have a circular chromosome, roughly 1.7 Mb long (Chen et al., 2013; Parkhill et al., 2000; Pearson et al., 2013; Taylor, Eaton, Yan, & Chang, 1992) which encodes for approximately 1,700 genes (Parkhill et al., 2000). A core genome multilocus sequence type (cgMLST) typing scheme has been defined jointly for these species which contains a set of 1,343 loci which are present in most (*∼*95%) members of human *C. jejuni* and *C. coli* isolates (Cody, Bray, Jolley, McCarthy, & Maidena, 2017). In any given dataset, an isolate will contain nearly all of these genes in this scheme, however the observed alleles of each gene are commonly found in only one or a few isolates. This means that there are many alleles across the genome which would be unique to individual collections of isolates from human and animal datasets.

#### 2.3.2 Dataset

The Source Assigned Campylobacteriosis in New Zealand Study (SACNZ) is a source-assigned case-control study of notified human cases of campylobacteriosis in the Auckland and MidCentral District Health Board regions, New Zealand, between 2018-2019 (Lake et al., 2021). *Campylobacter jejuni* and *C. coli* isolates were cultured from these human cases, as well as from poultry, sheep, and cattle processors serving the Auckland and MidCentral District Health Boards. Whole genome sequencing (WGS) was carried out on the study isolates, with the microbiology and WGS procedures being described elsewhere (Lake et al., 2021). Following sequencing, draft genomes were assembled using the nullarbor2 pipeline^6^ with default settings and cgMLST allele sequences were found by BLAST analyses (Altschul, Gish, Lipman, Miller, & Myers, 1990) against known alleles from the PubMLST *Campylobacter* database (Cody et al., 2017). Previously found and novel alleles were aligned using mafft (Katoh, Misawa, i. Kuma, & Miyata, 2002; Katoh & Standley, 2013) and an allele number assigned.^7^

The SACNZ dataset consists of 1,211 isolates from four sources: cattle (168), chicken (205), sheep (187), and human (651). Each isolate has an allelic profile consisting of the pattern of alleles across 1343 genes. The allelic designation for each gene identifies the unique aligned sequence for a previously described allele or a novel allele sequence. More simply, the categorical predictor variables are genes with alleles as levels. We use nucleotide sequencing information to calculate a matrix of Hamming distances (“Hamming Distance”, 2009) between each pair of alleles within each gene.

#### 2.3.3 Cross Validation

The 651 isolates collected from humans were excluded from analysis because their true animal source was unknown, and the remaining 560 isolates were subject to ten-fold cross-validation for each of three methods (CA, CA-unbiased, and PCO) using the same random number seed. Across the methods, the forest consisted of 500 trees and the “gini” index was used as the splitting criterion. For each method, ten independent random forest models were run (one on each of the ten folds) allowing each of the 560 isolates to be represented exactly once in testing data. Model performance was assessed by calculating the proportion of incorrect classifications on the set of test data for each fold and calculating the average and standard error, accounting for any variation between folds. Thus 560 isolates of known source were classified by a random forest model containing 500 trees resulting in 280,000 individual tree predictions for each method. To assess the effect of absent levels on classification success the number of absent levels used by each tree for prediction was recorded in addition to the individual tree predictions.

The order of analyses was as follows (see also figure 1):

1. create training and testing data
  - split the data into ten folds
  - select nine of the ten folds for a set of training data and the remaining tenth fold for a set of testing data
  - repeat until ten unique sets of training data and testing data have been created for each set and continue to 2.
2. prepare training data
  - create a level by class (i.e., allele by source) contingency table (CA-encoding, CA-unbiased-encoding methods)
  - encode each variable *via* principal component analysis (PCA) on the (weighted) contingency table (CA-encoding, CA-unbiased-encoding methods)
  - encode each variable *via* PCO on an ancillary set of data matched to the training data (PCO-encoding method)
3. fit the model on the prepared training data
4. prepare testing data
  - identify levels that are unique to the testing data (i.e., absent levels)
  - encode levels that are in the training data with the variable score from 2.
  - encode absent levels
    - with a score of infinity (CA-encoding method);
    - with a score of zero (CA-unbiased-encoding method)
    - with new scores *via* Gower’s method (Gower, 1968) on ancillary data matched to the testing data (PCO-encoding method)
5. predict each test observation
  - identify individual tree predictions
  - identify trees that branched on an absent level

#### 2.3.4 Code Availability

All analyses were carried out using R version 4.3.0 (R Core Team, 2023) and the ranger package (“RANdom forest GEneRator”) (Wright & Ziegler, 2017). The R code used in this study is available at https://github.com/smithhelen/LostInTheForest.

## 3 Results

### 3.1 Genome Description

Of the 560 isolates, there were 558 distinct allelic profiles (i.e., only 2 isolates shared an identical set of alleles with another isolate and the remaining isolates differed by at least one allele across the core genome). The number of alleles per gene ranged from 1 to 222 (median 35) and the total number of alleles was 49,424. Across all 1,343 genes, 25,317 alleles (51.2%) were seen in only a single source, and 17,575 alleles (35.6%) were seen in only a single isolate. 167/168 (99.4%) of the cattle isolates, 204/205 (99.5%) of the chicken isolates, and 187/187 (100%) of the sheep isolates contained alleles unique to their respective source. The unaligned sequence length of the genes ranged from 95 to 4,554 nucleotides (median 816). The number of nucleotides that differed between any pair of alleles (the Hamming distance) in aligned sequences ranged from 1 to 2,595 (median 42).

### 3.2 Random Forest Results

At least 90% of the random forest predictions, from any method, used at least one absent level for classification, and approximately one fifth (16% (PCO); 22% (CA and CA-unbiased)) of individual tree predictions used at least one absent level. The frequency of absent level use in predictions varied considerably among individual trees and forests for all methods. The CA-unbiased-encoding methods used absent levels up to 24 times in a single tree, compared with 19 for the PCO-encoding method and 12 for the CA-encoding method. On average, a variable with absent levels was used for a single classification between 4.7 times (PCO) and 7.5 times (CA-unbiased) but fewer than 4% of trees, from any method, used a variable with absent levels more than once for a single tree prediction.

The ten most important predictor variables (genes) as measured by the permutation variable importance approach (Breiman, 2001) varied between methods. CA-encoding and CA-unbiased-encoding methods identified the same 10 genes, in identical order. Of these ten only one was identified by the PCO-encoding method.

### 3.3 Classification Accuracy

The CA-unbiased-encoding method had the lowest average misclassification error (23.2%*±*1.2%), followed by the PCO-encoding (25.2%*±*1.2%), and the CA-encoding (27.0%*±*1.2%) methods. The accuracy of predictions was dependent on the class being predicted (table 1, figure 2). Across the methods, isolates sourced from chicken were the most accurately classified (79.7% *±*2.1% *−*84.2%*±*2.0%); isolates that were incorrectly classified were evenly distributed between sheep and cattle. Isolates sourced from sheep were the second most accurately classified for all methods (77.2% *±*2.0% *−*78.4% *±*2.1%); incorrectly classified isolates were mostly assigned to cattle (16.6% *±*1.9% *−*17.9% *±*2.0%) with fewer than 8% being assigned to chicken. Isolates sourced from cattle had the lowest classification success rates (60.5% *±* 2.1% *−* 68.4% *±* 2.1%), with most of the incorrect classifications predicted as sheep (20.8% *±* 2.2% *−* 31.0% *±* 2.2%) rather than chicken (10.9% *±* 4.4% *−* 13.4% *±* 4.6%).

**Table 1.**
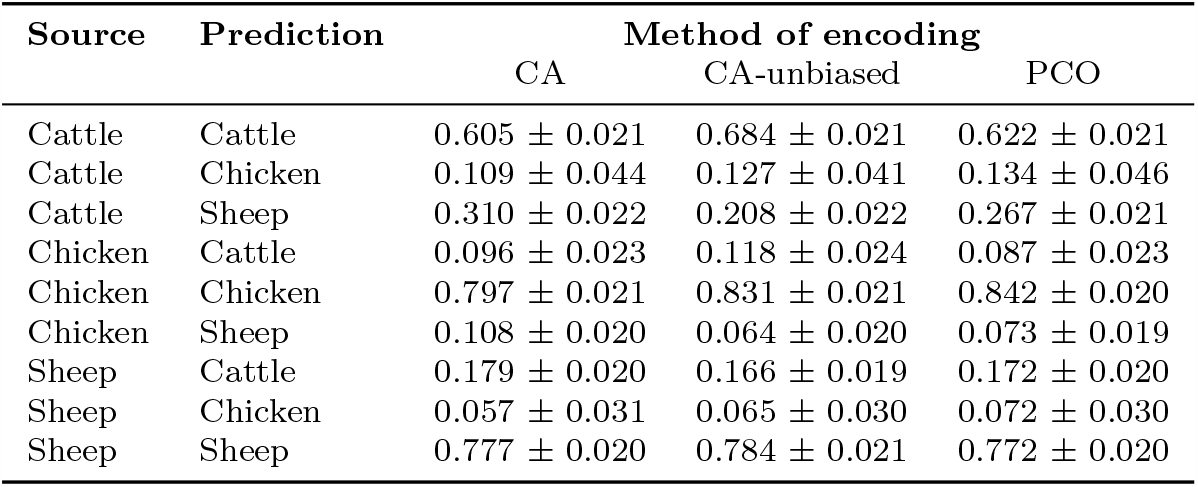
Weighted average proportion and standard error of all tree predictions assigned to each of three host sources (cattle, chicken and sheep) for each of three methods of encoding categorical predictors.

**Fig. 2.**
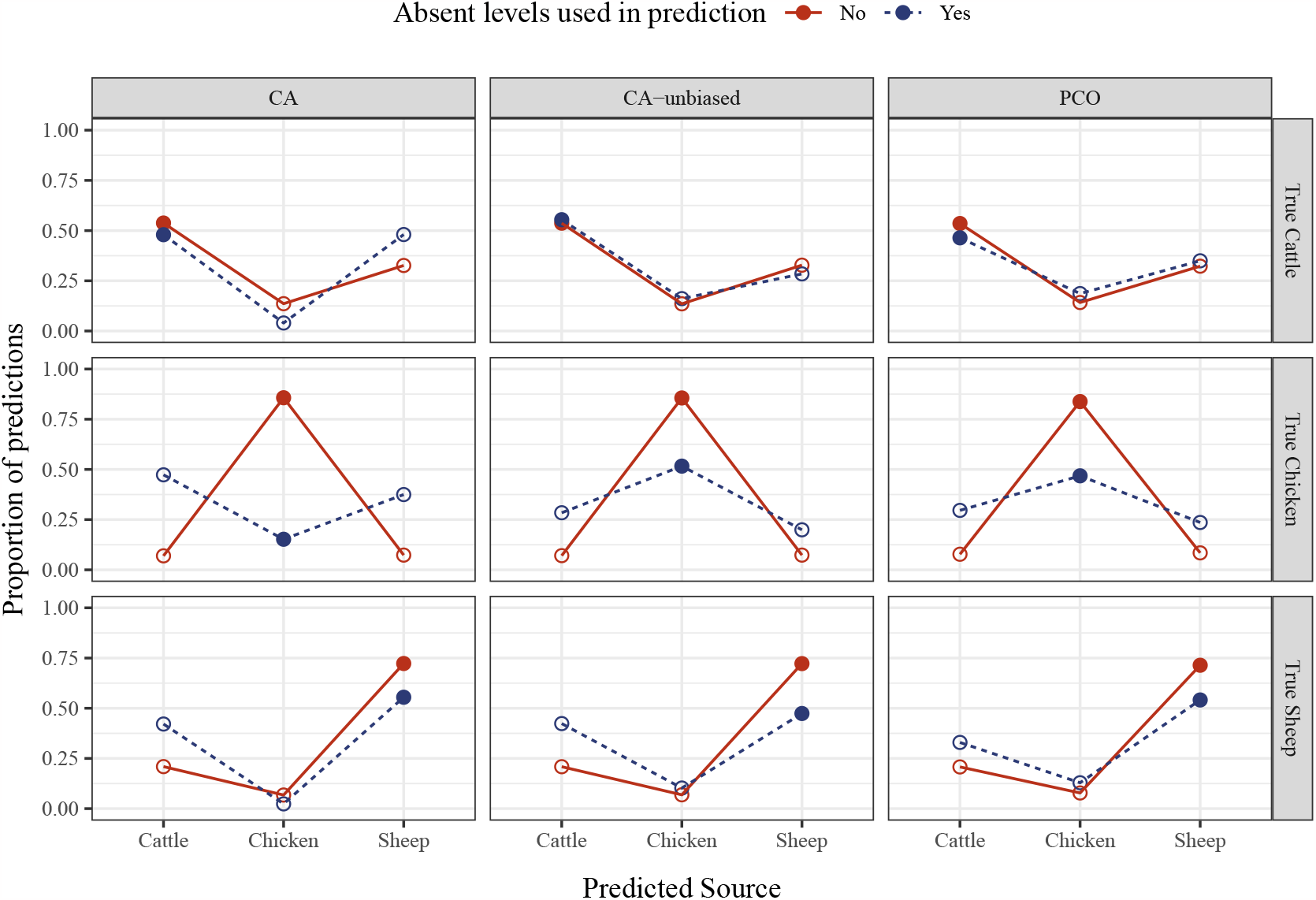
Proportion of tree predictions assigned to each of three host sources (cattle, chicken and sheep) when absent levels are used or not used in predictions. Open circles represent the proportion of cases for which the true class is predicted incorrectly; closed circles represent the proportion of cases for which the true class is predicted correctly.

### 3.4 Effect of Absent Levels

The class frequencies of predictions were similar across all methods when no absent levels were used for the predictions (figure 2). When absent levels were used for predictions, the predictions were not equally distributed across the three sources and the pattern of distribution depended on the method. For all methods, the class distribution followed the pattern of distribution for predictions made without absent levels, whereby incorrect chicken predictions were split between cattle and sheep; incorrect sheep classifications favoured cattle; and incorrect cattle classifications favoured sheep, but with a lower proportion of correct predictions in any class (figure 2). The accuracy of predictions also decreased as the number of absent levels in a tree increased (figure 3, see also figure S2).

**Fig. 3.**
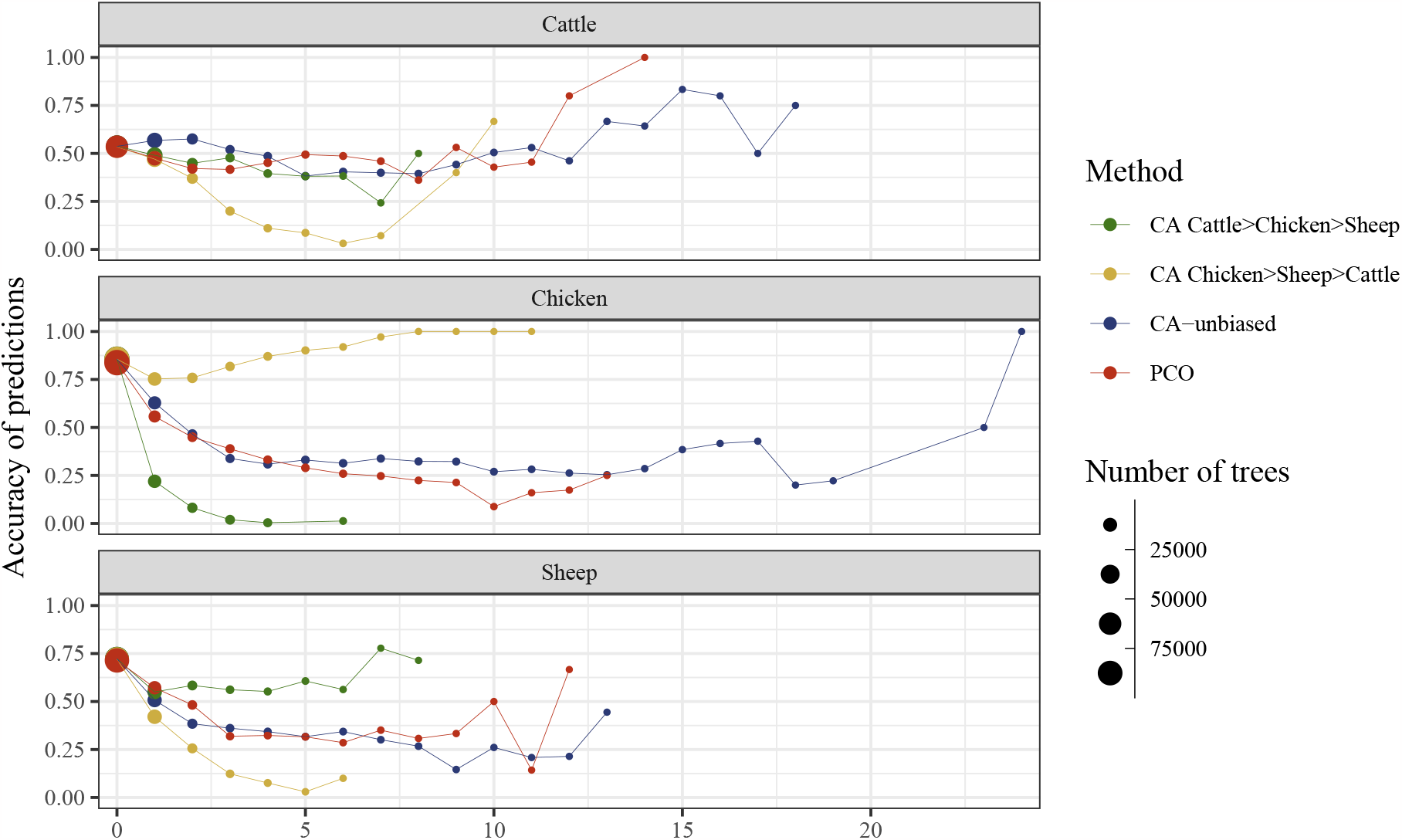
Proportion of predictions which were correct for trees with different numbers of absent levels and different methods and/or ordering of response class.

### 3.5 Effect of Response Class (Source) Order

The order of the response (source) levels also affected the success rates of predictions for the CA-encoding method when absent levels were used in prediction (figure 4). By default, most software treats the levels of categorical variables alphabetically, unless another ordering is specified explicitly. For our data this equates to cattle *<* chicken *<* sheep. In the presence of absent levels, the CA-encoding method will encode any absent level with the highest rank and thus the observations will always be sent down the right branch of the tree. When the source levels were re-ordered as chicken *<* sheep *<* cattle, more observations with an absent level were assigned to chicken (the first response) than when the default ordering was used. This effect of class order did not occur with the CA-unbiased-encoding, or PCO-encoding methods (figure S1).

**Fig. 4.**
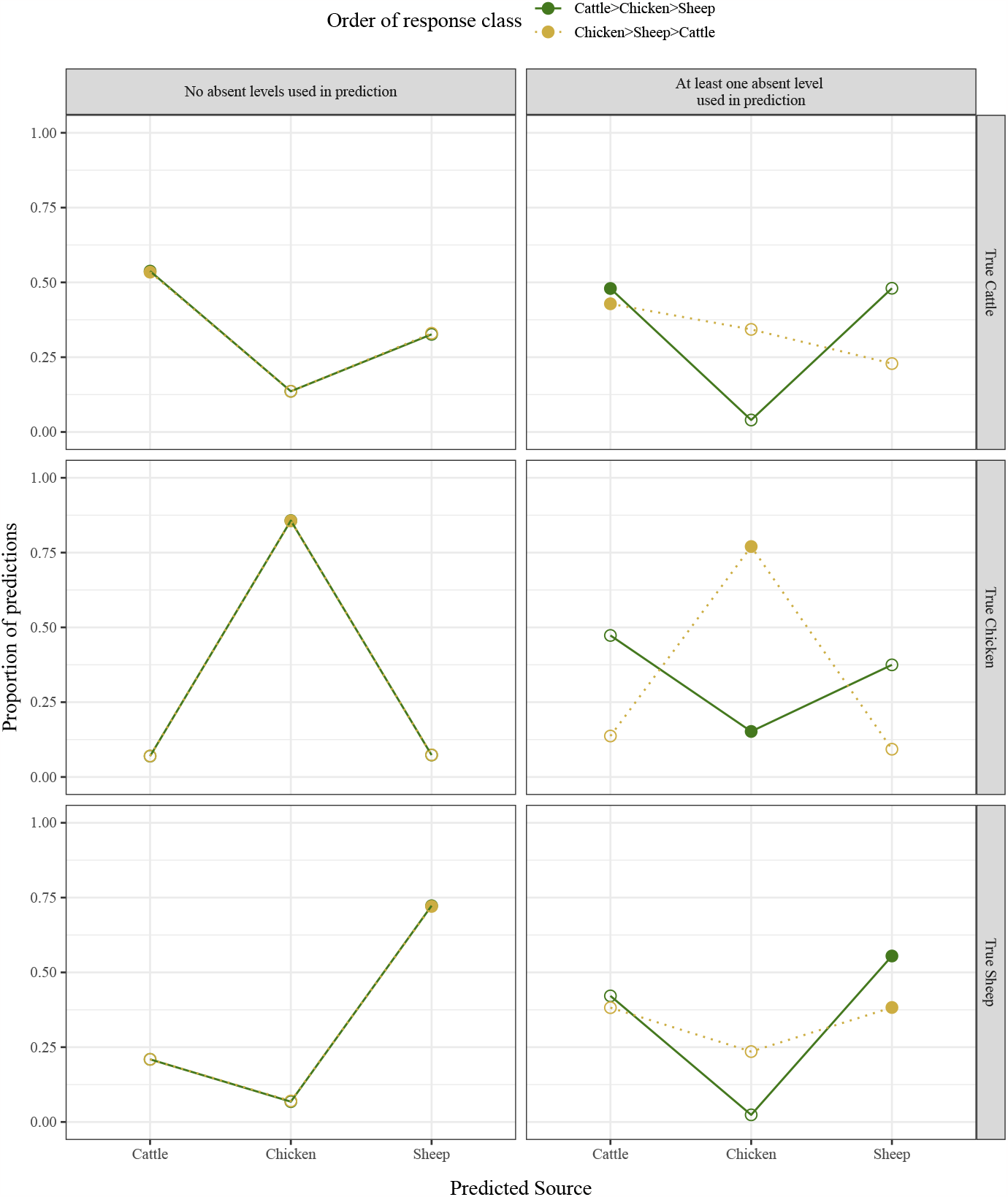
The effect of response class order on classification accuracy for the CA-encoding method. Open circles represent the proportion of cases for which the true class is predicted incorrectly; closed circles represent the proportion of cases for which the true class is predicted correctly.

## Discussion

Data sets with large numbers of predictor variables and/or large numbers of categories create a significant challenge for modelling. Random forest is a compelling option for such cases, particularly suited to sets of high dimensional data of high cardinality. Random forest models trained with high cardinality variables, such as source attribution models utilising a core genome MLST scheme, will almost certainly encounter absent levels when predicting for new data, and indicator encoding would lead to a prohibitively large number of binary variables - the cumulative number of unique alleles, across the genomes, from all the observations used to train the model.

Ordinal encoding can result in significant gains in efficiency of random forest models and additionally bypasses any restrictions imposed on the number of levels^8^. Ordinal encoding also provides a means of classifying observations with absent levels as additional levels can be added sequentially. The ordinality induced by integer encoding is artificial, however, and may be detrimental to random forest predictions (Wright & König, 2019). It is particularly problematic if the alphabetical ordering of the levels (i.e., the labelling) has some degree of association with the class, which may occur with temporal labelling of predictor levels. For example, the open-access PubMLST database (Jolley, Bray, & Maiden, 2018) defines alleles numerically and in a sequential manner based on sequence deposition. In this instance, treating alleles as numeric would not be appropriate because allele “1” is not necessarily more related to allele “2” than it is to allele “500”. However, it is likely that isolates have been added to the database in groups according to host source, so that their numeric order may partition into contiguous chunks by host. The numeric order thus provides information on likely host sources which is external to the data in a particular study, potentially biasing class assignment.

We found that, for random forests, different methods of encoding nominal variables had important implications for the accuracy of predictions when absent levels were encountered during prediction. When predicting using data with absent levels the CA-encoding method was biased towards the first response class. We also found that the systematic bias was affected by both the proportion of absent level in the data as well as the level of association of the absent level with a response class (S2). For this method, the predictor levels are target encoded using their contribution to response class and an absent level is encoded with the highest rank. Changing the order of levels of the response classes can alter (reverse) the ranks of the predictor levels, however, the absent level will always retain the highest rank. Thus, the absent level will be next in rank to a level of a predictor associated with one response class in one ordering, but with the reverse ordering it will be next in rank to a different predictor level, potentially associated with a different response class. This option for encoding variable levels has previously been recommended when variables have a large number of levels and/or do not have an inherent order (Wright & König, 2019).

Our first alternative method, the CA-unbiased-encoding method, is identical to the CA-encoding method except for the treatment of absent levels. The CA-unbiased-encoding method encodes all absent levels with a score of zero (rather than infinity) in line with the assumption of *a priori* equal class probabilities. This approach resolved the systematic bias towards the first response class caused by absent levels and showed a small improvement in overall classification accuracy (S2).

Our second alternative method, the PCO-encoding method, used Gower’s method of principal coordinates analysis on data that was independent of the class probabilities to inform the encoding of predictor variables, including absent levels (figure 1, c). This method assumes that an observation with an absent level is more likely to branch in the same direction as an observation whose corresponding level is ‘similar’ to the absent level. This requires information with which to quantify the similarity (or dissimilarity) of each pair of levels of a predictor variable. We demonstrated the method using genomic sequencing data for each predictor variable, more specifically, the number of nucleotides shared by any two alleles (Hamming distance) for a given gene. In contrast to the CA-encoding and CA-unbiased-encoding methods, encoding using PCO was independent of the counts of levels of predictor variables in the training data, and thus also able to be applied to absent levels. In addition, rather than encoding all absent levels with the same score, the PCO-encoding method encoded each absent level individually. Using the Hamming distance between the absent allele and every other allele, the absent allele was encoded so that it was more similar to an allele with which it shared more nucleotides and less similar to an allele with which it shared few nucleotides. This is based on the assumption that isolates from one source would be more likely to have alleles which are similar in terms of their genome sequence, than isolates from another source (Pinheiro, de Souza Pinheiro, & Sen, 2005; Pérez-Reche, Rotariu, Lopes, Forbes, & Strachan, 2020). This method was not systematically biased, and had similar prediction accuracy to the CA-unbiased-encoding method.

The issue with absent levels will be less problematic for data where every level of every predictor variable in the set of observations to be classified is present in the training data, and more problematic for data containing variables with many levels. Previously, it was thought that no meaningful splitting decision can be made for observations with new levels at a splitting node and discussion has ensued regarding the advantages of keeping the observations with absent levels together *versus* assigning them randomly at a split (Wright & König, 2019). We introduced two methods which do make meaningful splitting decisions for observations with new levels - the CA-unbiased-encoding method and the PCO-encoding method. Both of these methods produce competitive prediction results, resolve the systematic bias caused by absent levels, and avoid arbitrary splitting decisions for observations with absent levels. Although here only the first principal component/coordinate is used, it may be beneficial to increase the dimension to at least two principal components/coordinates. In addition, combining a target-based approach with ancillary information on the levels to inform variable ordering, particularly the placement of absent levels, may further improve classification success. These new methods would be suitable for any high cardinality predictors where a measure of level similarity could be determined. For example, a free text response field from a survey has a potentially infinite set of responses and absent levels would be almost inevitable. The string difference between responses could be used as a measure of similarity.

The success of a random forest classification model is often measured by the rate of misclassifications. Breiman (1996, 2001) claimed that the out-of-bag misclassification rate (i.e., the rate of misclassification of cases that were not selected for training a particular tree) was as reliable as using an independent set of data for testing. When using a target-based encoding method (e.g., the CA-encoding or CA-unbiased-encoding methods), there is information leakage from the target variable to the predictors. The levels of each predictor variable are encoded according to the first principal component of the weighted matrix of class probabilities, calculated from the entire (training) dataset before the analysis. Each observation in the set of training data is used to train approximately two thirds of the trees in the forest. The remaining third of trees can be used to generate an OOB prediction for that observation, which will be either correct or not. There is information leakage, however, because even when the observations are in the OOB set, the encoding of their corresponding levels was informed from the entire dataset (i.e., prior to the observations moving OOB) based on the correct response classes (i.e., the target); therefore, the OOB observations do not behave like fully independent test data. This leakage will impact OOB errors and they will likely underestimate the true misclassification rates. Potential solutions to this problem include re-ordering the levels at each split in the tree, re-ordering the levels of each bootstrap sample, or calculating the misclassification rate based on a fully independent test dataset. Target-agnostic encoding methods, such as the naïve alphabetical encoding and the PCO-encoding method, do not suffer the information-leakage problem because the response class (target) information is not used for the encoding. The PCO-encoding method will therefore not have this potential issue with incorrect OOB misclassification rates.

## 5 Conclusion

This paper highlights potential pitfalls in the use of classification trees when an order is imposed on nominal predictor variables. These findings are applicable to random forests and other tree-based methods (e.g., boosted trees) when new levels of categorical predictor variables are encountered during prediction and/or where OOB misclassification rates are calculated. When levels of categorical predictor variables are target encoded using class probability information, and absent levels are integrated at the highest rank (effecting a consistent direction for them to branch at a split), predictions were systematically biased to the first response class. Target-based encoding of predictors using class probability information, and integrating absent levels according to the *a priori* hypothesis of equal class probability, is a potential and unbiased solution with good predictive properties. Target-agnostic encoding of predictors using information which quantifies the similarity between each pair of predictor levels, and integrating absent levels by virtue of their similarity to each of the other levels in the training data, is another potential solution which removes the need for arbitrary decisions on where to direct absent levels. This approach has good predictive properties, is not biased, and does not affect the OOB misclassification rate. The predictive performance of the PCO-encoding method depends on the ability to separate the levels according to class in the principal co-ordinate space and will depend on the ancillary information available. For high cardinality data, such as WGS data, it is almost certain there will be absent levels across the predictor variables, and that a large number of observations will be affected. Removing observations and/or variables with absent levels is, therefore, not a viable option. When there are no, or few, absent levels the different methods have similar predictive performance. However, as there can never be assurance of an absence of absent levels, there are no circumstances where the CA-encoding method should be used. We recommend, when ancillary information is available, such as with WGS data, the PCO-encoding method for random forests and suggest comparing model performance with the CA-unbiased-encoding method using misclassification rates calculated with an independent dataset. A reduction in bias for source attribution modelling will lead to a better understanding of potential risk factors in zoonotic infectious diseases to better inform public health decision making.

## Supplementary information

### S1 The effect of response class order on classification accuracy

**Fig. S1.**
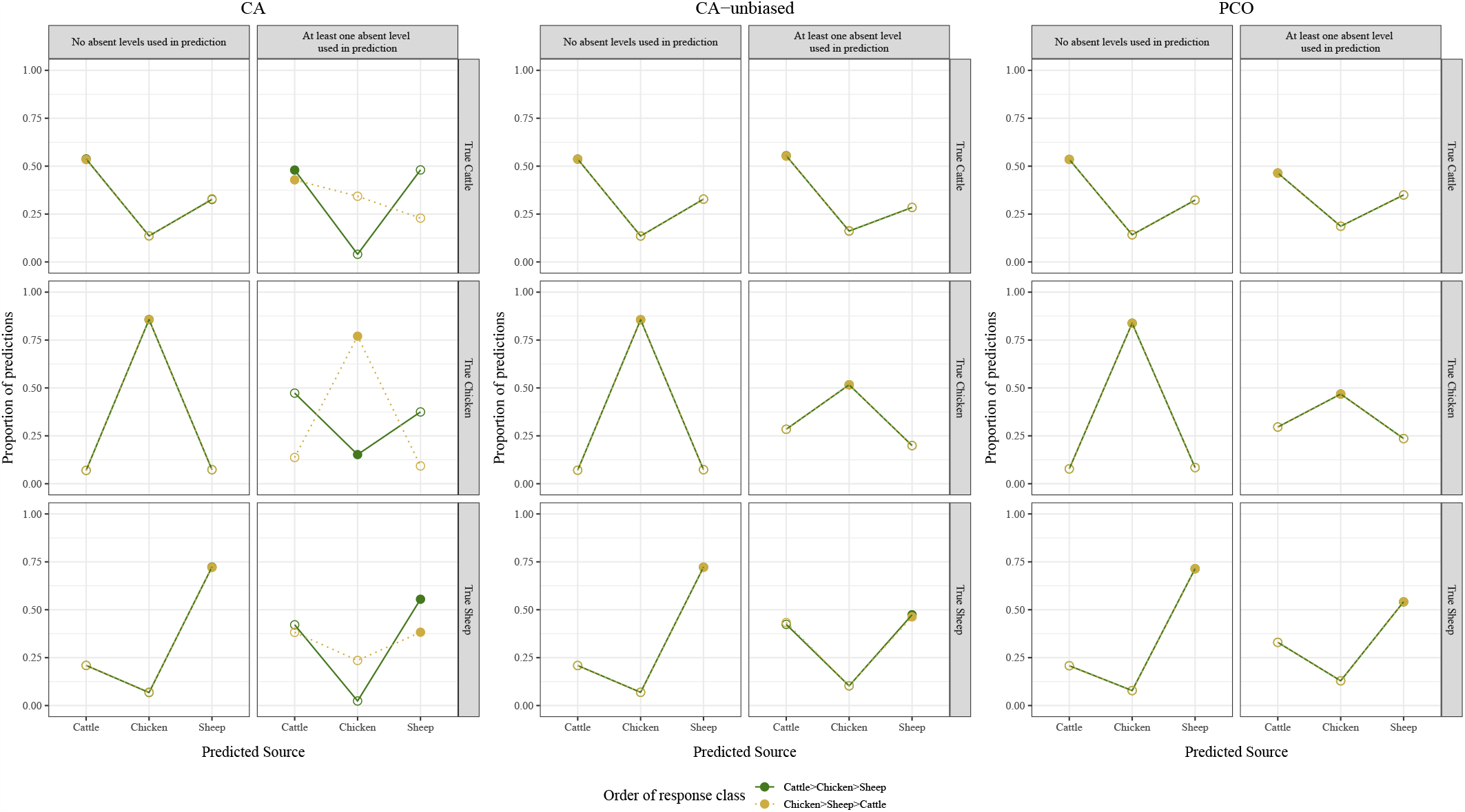
The effect of response class order on classification accuracy. Open circles represent the proportion of cases for which the true class is predicted incorrectly; closed circles represent the proportion of cases for which the true class is predicted correctly.

This article has two accompanying supplementary files -

S1 The effect of response class order on classification accuracy;

S2 Bias resulting from treatment of absent levels

### S2 Bias resulting from treatment of absent levels

**Fig. S2.**
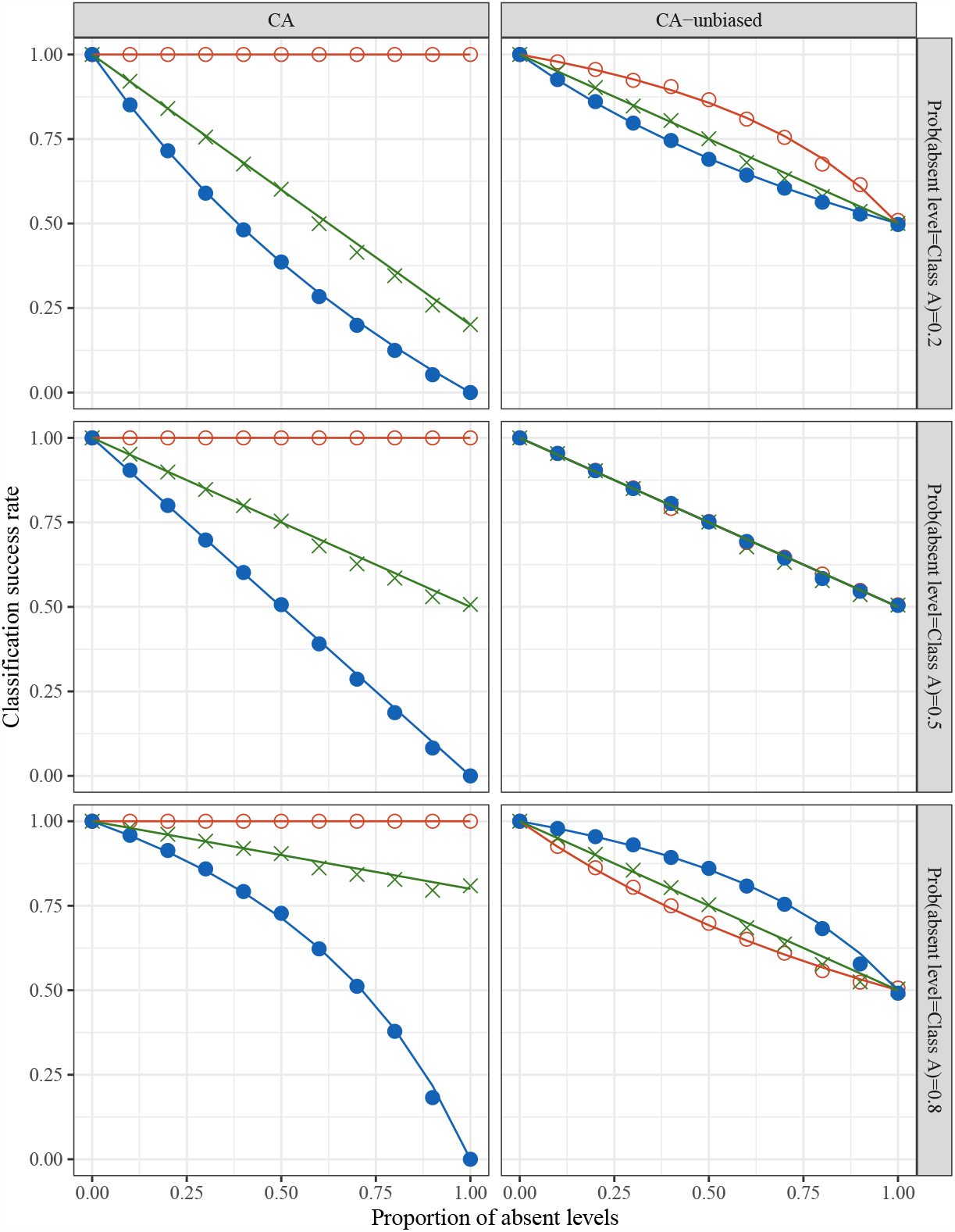
The effect of absent levels on classification accuracy for the CA-encoding and CA-unbiased-encoding methods. Open circles represent the proportion of cases for which Class A is predicted correctly; closed circles represent the proportion of cases for which Class B is predicted correctly; crosses represent the proportion of total cases which were predicted correctly; the lines represent the expected probabilities.

To examine how the bias resulting from treatment of absent levels is affected by the proportion of absent levels, a set of data was simulated and analysed with random forest following encoding *via* the CA-encoding and the CA-unbiased-encoding methods. The simulated dataset consisted of

n observations coming from one of two classes (A and B), each with a single predictor variable that took one of 3 levels (a, b, or c). Levels a and b were perfectly associated with class A and B respectively, while level c was associated with class A with probability *p*_*A*_ which varied from *p*_*A*_ *∈* 0.2, 0.5, 0.8.

The training set consisted of the predictor with levels a and b leading to perfect separation; the test set consisted of the absent level c with probability *p*_*c*_ *∈ {*0, 0.2, 0.4, 0.6, 0.8, 1}, with levels a and b assigned with probability (1− *p*_*c*_)*/*2.

In this simple scenario we can assess the missclassification rate exactly. Let m be the probability of the observation being sent to the right branch of the split, where

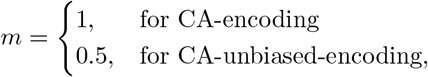

then the expected misclassification rate of class A is

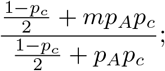

the expected misclassification rate of class B is

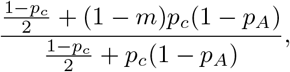

and the expected total misclassification rate is

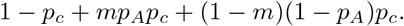

We found that the simulations matched the expected probabilities (figure S2). When absent levels are scored as infinity, as per the CA-encoding method, the random forest model is biased towards the first response class (Class A) and this bias gets worse with increasing proportion of absent levels and increasing association of absent level with class B. In addition, the overall misclassification rate for the CA-encoding method is affected by the level of association of the absent level with the response class. When absent levels are scored as zero, as per the CA-unbiased-encoding method, the random forest model favours the response class with the greatest association with the absent level but it is not affecting the overall misclassification rate which is independent of the level of association of the absent level with either response class.

Different methods of encoding nominal variables have important implications for the accuracy of error rates when absent levels are present in the data. The CA-encoding method is biased when classifying observations with absent levels. When there are no, or few, absent levels both methods have similar predictive performance, however the CA-encoding method never out-performs the CA-unbiased-encoding method.

## Acknowledgments

This research is supported by a Massey University School of Fundamental Sciences scholarship.

## Footnotes

1 https://scikit-learn.org/stable/modules/ensemble.html#random-forests

2 https://github.com/imbs-hl/ranger/issues/94

3 Coppersmith, Hong & Hosking (1999) use the first principal component of the weighted matrix of class probabilities

4 Update 4.6-10 allows absent levels to be encoded if the categorical variable is an ordered factor. Categorical variables of type “character” are converted to ordered factors, with the order determined alphabetically.

5 The “ordered” method in `ranger` performs a PCA on the weighted covariance matrix of class probabilities rather than on the weighted matrix of class probabilities, yet the results are equivalent.

6 https://github.com/tseemann/nullarbor

7 https://github.com/jmarshallnz/cgmlst

8 When nominal encoding a categorical variable (e.g. the “`partition`” method in `ranger`), each binary node assignment is saved using the bit representation of a double integer, which limits this treatment to predictors with fewer than 54 levels (Wright & König, 2019).

## References

Altschul, S.F., Gish, W., Lipman, D.J., Miller, W., Myers, E.W. (1990). Basic local alignment search tool [Journal Article]. Journal of Molecular Biology, 215 (3), 403–410,

Amit, Y., & Geman, D. (1997). Shape quantization and recognition with randomized trees [Journal Article]. Neural Computation, 9 (7), 1545–1588,

Arning, N., Sheppard, S., Bayliss, S., Clifton, D., Wilson, D. (2021). Machine learning to predict the source of campylobacteriosis using whole genome data [Journal Article]. PLoS Genetics, 17 (10),, 10.1371/journal.pgen.1009436

Au, T.C. (2018). Random forests, decision trees, and categorical predictors: The “absent levels” problem [Journal Article]. Journal of Machine Learning Research, 19, 1–30,

Breiman, L. (1996). Bagging predictors [Journal Article]. Machine Learning, 24 (2), 123–140,

Breiman, L. (2001). Random forests [Journal Article]. Machine Learning, 45 (1), 5–32,

Breiman, L., Friedman, J.H., Olshen, R.A., Stone, C.J. (1984). Classification and regression trees [Book]. Wadsworth International Group.

Brinch, M., Hald, T., Henri, C., Wainaina, L., Merlotti, A., Remondini, D., Njage, P. (2023). Comparison of source attribution methodologies for human campylobacteriosis. [Journal Article]. Pathogens, 12 (6)

Cerda, P., Varoquaux, G., Kegl, B. (2018). Similarity encoding for learning with dirty categorical variables [Journal Article]. Machine Learning, 107, 1477–1494,

Chen, Y., Mukherjee, S., Hoffmann, M., Kotewicz, M.L., Young, S., Abbott, J., … Zhao, S. (2013). Whole-genome sequencing of gentamicin-resistant campylobacter coli isolated from u.s. retail meats reveals novel plasmid-mediated aminoglycoside resistance genes [Journal Article]. Antimicrob Agents Chemother, 57 (11), 5398–5405,

Cody, A.J., Bray, J.E., Jolley, K.A., McCarthy, N.D., Maidena, M.C.J. (2017). Core genome multilocus sequence typing scheme for stable, comparative analyses of campylobacter jejuni and c. coli human disease isolates [Journal Article]. Journal of Clinical Microbiology, 55 (7), 2086–2097,

Coppersmith, D., Hong, S.E.J., Hosking, J.R.M. (1999). Partitioning nominal attributes in decision trees [Journal Article]. Data Mining and Knowledge Discovery, 3 (2), 197–217,

Domingues, A.R., Pires, S.M., Halasa, T., Hald, T. (2012). Source attribution of human campylobacteriosis using a meta-analysis of case-control studies of sporadic infections [Journal Article]. Epidemiol Infect, 140 (6), 970–981,

Fisher, W.D. (1958). On grouping for maximum homogeneity [Journal Article]. Journal of the American Statistical Association, 53 (284), 789–798,

Gower, J.C. (1966). Some distance properties of latent root and vector methods used in multivariate analysis [Journal Article]. Biometrika, 53 (3/4), 325–338,

Gower, J.C. (1968). Adding a point to vector diagrams in multivariate analysis [Journal Article]. Biometrika, 55 (3), 582–585,

Hald, T., Aspinall, W., Devleesschauwer, B., Cooke, R., Corrigan, T., Havelaar, A.H., … Hoffmann, S. (2016). World health organization estimates of the relative contributions of food to the burden of disease due to selected foodborne hazards: A structured expert elicitation [Journal Article]. PLoS One, 11 (1), e0145839,

Hald, T., Vose, D., Wegener, H.C., Koupeev, T. (2004). A bayesian approach to quantify the contribution of animal-food sources to human salmonellosis [Journal Article]. Risk Anal, 24 (1), 255–269,

Hamming distance. (2009). In S. Li & A. Jain (Eds.), Encyclopedia of biometrics (pp. 668–668). Boston, MA: Springer US.

Hastie, T., Tibshirani, R., Friedman, J.H. (2009). The elements of statistical learning : data mining, inference, and prediction [Book]. Springer.

Havelaar, A.H., Galindo, A.V., Kurowicka, D., Cooke, R.M. (2008). Attribution of foodborne pathogens using structured expert elicitation [Journal Article]. Foodborne Pathog Dis, 5 (5), 649–659,

Ho, T.K. (1998). The random subspace method for constructing decision forests [Journal Article]. IEEE Transactions on Pattern Analysis and Machine Intelligence, 20 (8), 832–844,

Hothorn, T., & Zeileis, A. (2015). partykit: A modular toolkit for recursive partytioning in r. Journal of Machine Learning Research, 16 (118), 3905–3909,

Ishwaran, H., Kogalur, U.B., Blackstone, E.H., Lauer, M.S. (2008). Random survival forests [Journal Article]. The Annals of Applied Statistics, 2 (3), 841–860,

Janitza, S., & Hornung, R. (2018). On the overestimation of random forest’s out-of-bag error [Journal Article]. PLoS ONE, 13 (8), e0201904,

Jolley, K.A., Bray, J.E., Maiden, M. (2018). Open-access bacterial population genomics: Bigsdb software, the pubmlst.org website and their applications [version 1; referees: 2 approved] [Journal Article]. Wellcome Open Research, 3, 124,

Katoh, K., Misawa, K., i. Kuma, K., Miyata, T. (2002). Mafft: a novel method for rapid multiple sequence alignment based on fast fourier transform [Journal Article]. Nucleic acids research, 30 (14), 3059–3066,

Katoh, K., & Standley, D.M. (2013). Mafft multiple sequence alignment software version 7: Improvements in performance and usability [Journal Article]. Molecular Biology & Evolution, 30 (4), 772–780,

Lake, R.J., Campbell, D.M., Hathaway, S.C., Ashmore, E., Cressey, P.J., Horn, B.J., … French, N.P. (2021). Source attributed case-control study of campylobacteriosis in new zealand [Journal Article]. International Journal of Infectious Diseases, 103, 268–277,

Liao, S.J., Marshall, J., Hazelton, M.L., French, N.P. (2019). Extending statistical models for source attribution of zoonotic diseases: a study of campylobacteriosis [Journal Article]. J R Soc Interface, 16 (150), 20180534,

Liaw, A., & Wiener, M. (2002). Classification and regression by randomforest. R News, 2 (3), 18–22, Retrieved from https://CRAN.R-project.org/doc/Rnews/

Miller, P., Marshall, J., French, N., Jewell, C. (2017). sourcer: Classification and source attribution of infectious agents among heterogeneous populations [Journal Article]. PLoS Comput Biol, 13 (5), e1005564,

Mitchell, M.W. (2011). Bias of the random forest out-of-bag (oob) error for certain input parameters [Journal Article]. Open Journal of Statistics, 01 (03), 205–211,

Müllner, P., Jones, G., Noble, A., Spencer, S.E., Hathaway, S., French, N.P. (2009). Source attribution of food-borne zoonoses in new zealand: a modified hald model [Journal Article]. Risk Anal, 29 (7), 970–984,

Parkhill, J., Wren, B.W., Mungall, K., Ketley, J.M., Churcher, C., Basham, D., … Barrell, B.G. (2000). The genome sequence of the food-borne pathogen campylobacter jejuni reveals hypervariable sequences [Journal Article]. Nature, 403 (6770), 665–668,

Pearson, B.M., Rokney, A., Crossman, L.C., Miller, W.G., Wain, J., van Vliet, A.H. (2013). Complete genome sequence of the campylobacter coli clinical isolate 15-537360 [Journal Article]. Genome Announc, 1 (6), e01056–13,

Pedregosa, F., Varoquaux, G., Gramfort, A., Michel, V., Thirion, B., Grisel, O., … Duchesnay, E. (2011). Scikit-learn: Machine learning in python [Journal Article]. Journal of Machine Learning Research, 12, 2825–2830,

Pinheiro, H.P., de Souza Pinheiro, A., Sen, P.K. (2005). Comparison of genomic sequences using the hamming distance [Journal Article]. Journal of Statistical Planning and Inference, 130 (1), 325–339,

Pintar, K.D.M., Thomas, K.M., Christidis, T., Otten, A., Nesbitt, A., Marshall, B., … Ravel, A. (2017). A comparative exposure assessment of campylobacter in ontario, canada [Journal Article]. Risk Anal, 37 (4), 677–715,

Pires, S.M., Vigre, H., Makela, P., Hald, T. (2010). Using outbreak data for source attribution of human salmonellosis and campylobacteriosis in europe [Journal Article]. Foodborne Pathog Dis, 7 (11), 1351–1361,

Pérez-Reche, F.J., Rotariu, O., Lopes, B.S., Forbes, K.J., Strachan, N.J.C. (2020). Mining whole genome sequence data to efficiently attribute individuals to source populations [Journal Article]. Sci Rep, 10 (1), 12124,

Quinlan, J.R. (1993). C4.5 : programs for machine learning. [Book]. Morgan Kaufmann Publishers.

R Core Team (2023). R: A language and environment for statistical computing [Computer software manual]. Vienna, Austria. Retrieved from https://www.R-project.org/

Reilly, D., Taylor, M., Fergus, P., Chalmers, C., Thompson, S. (2022). The categorical data conundrum: Heuristics for classification problems - a case study on domestic fire injuries [Journal Article]. IEEE Access, 10, 70113–70125,

Saar-Tsechansky, M., & Provost, F. (2007). Handling missing values when applying classification models. Journal of Machine Learning Research, 8, 1625–1657,

Sheppard, S.K., Dallas, J.F., Strachan, N.J., MacRae, M., McCarthy, N.D., Wilson, D.J., … Forbes, K.J. (2009). Campylobacter genotyping to determine the source of human infection [Journal Article]. Clin Infect Dis, 48 (8), 1072–1078,

Sheppard, S.K., & Maiden, M.C. (2015). The evolution of campylobacter jejuni and campylobacter coli [Journal Article]. Cold Spring Harb Perspect Biol, 7 (8), a018119,

Strachan, N.J., Gormley, F.J., Rotariu, O., Ogden, I.D., Miller, G., Dunn, G.M., … Forbes, K.J. (2009). Attribution of campylobacter infections in northeast scotland to specific sources by use of multilocus sequence typing [Journal Article]. J Infect Dis, 199 (8), 1205–1208,

Taylor, D.E., Eaton, M., Yan, W., Chang, N. (1992). Genome maps of campylobacter jejuni and campylobacter coli [Journal Article]. J Bacteriol, 174 (7), 2332–7,

Therneau, T., Atkinson, B., Ripley, B. (2022). rpart: Recursive partitioning and regression trees [Computer Program]. Retrieved from http://CRAN.R-project.org/package=rpart

Wright, M.N., & König, I.R. (2019). Splitting on categorical predictors in random forests [Journal Article]. PeerJ, 2019 (2), e6339,

Wright, M.N., & Ziegler, A. (2017). Ranger: A fast implementation of random forests for high dimensional data in c++ and r [Journal Article]. Journal of Statistical Software, 77 (1), 1–17,

